# Native entanglement misfolding contributes to age-associated structural changes across the *Saccharomyces cerevisiae* proteome

**DOI:** 10.64898/2026.04.15.717356

**Authors:** Quyen V. Vu, Ian Sitarik, Daniel A. Nissley, Edward P. O’Brien

## Abstract

Aging at the subcellular level involves the simultaneous decline in the cell’s ability to maintain protein homeostasis and rise in misfolded proteins through a positive feedback loop. Here, we test if a widespread class of protein misfolding could contribute to proteome aging by examining if statistical associations exist between age-related changes in protein structure, measured by limited proteolysis mass spectrometry data of the aging *Saccharomyces cerevisiae* proteome, with structural annotations and molecular simulations. We find that globular proteins that are likely to exhibit entanglement misfolding are 121% more likely to exhibit age-related structural changes, and these changes are 59% more likely to be localized to natively entangled regions. Proteins containing native entanglements are seven-fold more likely to misfold, according to simulations, and populate long-lived, near-native misfolded states. Thus, the age-related structural changes in yeast proteins can be explained in part by the accumulation of misfolded proteins involving entanglements.

## Introduction

Aging is the progressive and systemic decline in subcellular and cellular functionality spanning all length scales and diverse biological processes^1–6^. Over time, cells accumulate molecular damage and experience chronic stress, giving rise to hallmark features of aging including genomic instability, epigenetic dysregulation, metabolic and mitochondrial dysfunction^7^, and a loss of protein homeostasis – referred to as proteostasis^8^. The proteostasis network, comprising protein synthesis, molecular chaperones, the ubiquitin–proteasome system, autophagic pathways, and functional deaggregases, loses efficiency with age. These inefficiencies lead to accumulation of misfolded and aggregation-prone proteins that are either nonfunctional or gain new, detrimental functions^9–12^. This loss of protein quality control compromises cellular viability and contributes to age-associated diseases^13^.

Positive feedback loops contribute to these proteostasis inefficiencies^14^. Misfolded proteins can sequester chaperones, rendering them non-functional, and thereby reduce the capacity of the remaining chaperones to refold proteins^15^. As more chaperones are sequestered the concentration of free misfolded proteins increases^9,10,14^. This increased concentration of misfolded proteins, which are more likely to aggregate, can overwhelm the capacity of functional deaggregases to resolubilize the proteins that agglomerate into insoluble aggregates, leading to runaway aggregation and more cell stress^10^. Age-related loss-of-function in key proteins influencing autophagy and proteasomal degradation further impact the cells’ ability to clear misfolded or chemically damaged proteins^16^. Regardless of whether these feedback loops reflect non-linear, threshold-driven failures of proteostasis - as described by the collapse model^9–11^ - or gradual, stochastic erosion of protein quality – as in the network drift model^12^ – the common element to both is protein misfolding and subsequent loss or gain of some new function. For this reason, any newly discovered mechanism of protein misfolding could be a yet undiscovered mechanism of cellular aging.

It is therefore noteworthy that a series of papers have recently provided evidence that there exists a previously unrecognized, wide-spread class of protein misfolding that has qualities relevant to aging^17–23^. This class of misfolding can form long-lived^21^, soluble native-like states that are non-functional, providing an opportunity for them to persist in cells for extended periods and contribute to an overall loss-of-function of the proteome^17^. Further, because some of these states are structurally similar to their protein’s native state – exhibiting localized misfolding along backbone segments – they have the potential to bypass the refolding action of chaperones^17,22^. Bypassing these proteostasis quality controls could lead to their accumulation during aging^10,24^. Both of these qualities have the potential to lead to increasing proteostasis inefficiencies with age^12^.

This wide-spread class of misfolding involves the geometric motif known as a non-covalent lasso entanglement (NCLE), in which a segment of the polypeptide backbone forms a loop, closed by a one or more pairs of residues interacting non-covalently, and through which another portion of the protein backbone is threaded (Fig. 1a-b). Simulation studies have shown changes of NCLE status can act as kinetic traps during protein folding^19,21^, resulting in long-lived misfolded states. Misfolding involving a change of entanglement occurs via three mechanisms^21,22^ (Fig. 1c): a gain of non-native entanglement (i.e., formation of a NCLE that does not exist in the native state), loss of a native entanglement (i.e., failure to form an NCLE that is present in the native fold), or some combination thereof^22^. Proteins without entanglements in their native state can misfold only through the former, whereas natively entangled proteins are susceptible to all three mechanisms. This expanded vulnerability means that native entangled proteins inherently increase a protein’s misfolding propensity^22^.

**Fig. 1:**
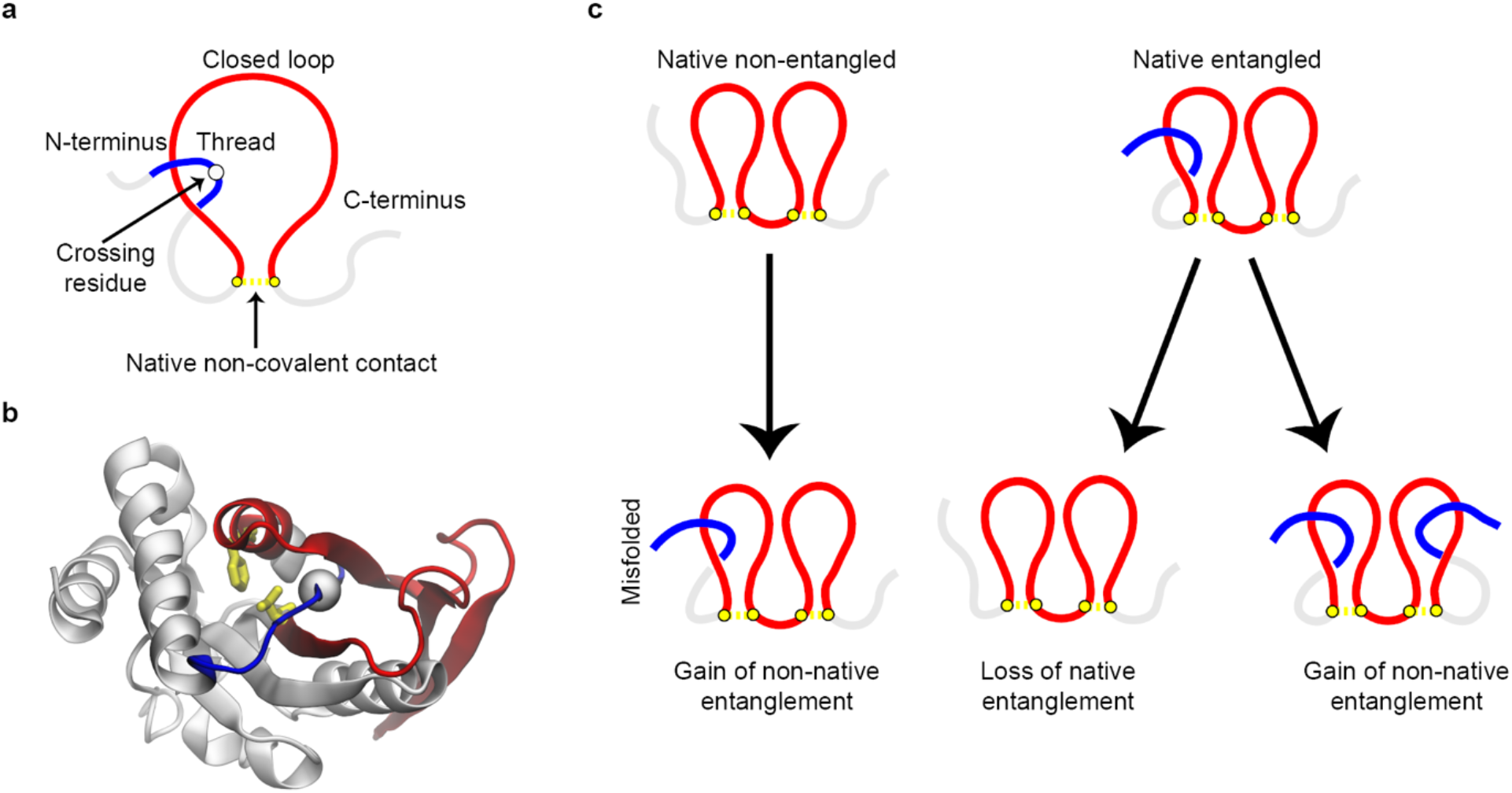
Non-covalent lasso entanglement (NCLE) and misfolding mechanisms that alter entanglement state. (a) Schematic illustration of the geometric elements that defined an NCLE. A loop (red) is closed by non-covalent contact (yellow) and is threaded by another segment (blue) at the crossing residue (white sphere). (b) Native structure of adenine phosphoribosyltransferase 1 (Uniprot: P49435), highlighting a native NCLE. The loop is shown in red and is closed by a native non-covalent contact between residue PHE73 and VAL128 (yellow licorice). The N-terminal threading segment is shown in blue and passes through the loop at the crossing residue ILE63 (white sphere). (c) Schematic of possible NCLE misfolding mechanisms for proteins without a native NCLE, left, and for proteins with native NCLEs, right.

We thus hypothesized that proteins containing native lasso entanglements are more susceptible to age-associated structural changes than non-entangled proteins. To test this hypothesis, here, we integrate structural entanglement annotations^25^ with a proteome-wide limited proteolysis (LiP) mass spectrometry (MS) dataset that measures across replicative age conformational changes in hundreds of proteins from *Saccharomyces cerevisiae* (*S. cerevisiae*)^26^. This is done by comparing the changes in polypeptide digestion patterns – which report on conformational changes - across the proteome of young (average age 0.6 ± 0.91 divisions) versus aged (average age 4.2 ± 0.65 divisions) cells. We also use molecular simulations of protein folding to gauge and correlate misfolding propensity of a diverse set of proteins. Our findings indicate that the misfolding of native NCLEs explain a large proportion of the conformational changes measured during aging. This suggests that NCLE misfolding could contribute broadly to the decline of proteome integrity and cellular function with age.

## Results

### Overview of the LiP-MS data

We analyzed a previously published LiP-MS dataset of young cells, having an average age of 0.6 ± 0.91 divisions, and aged cells, having an average age 4.2 ± 0.65 divisions, in *S. cerevisiae*^26^. In these mass spectrum data there are 2,833 detected proteins common across all conditions, and 2,292 proteins have high quality AlphaFold structures^27^ (average pLDDT ≥ 70). Amongst these high-quality structures, 1,706 have one or more native NCLEs, 21 have covalent lassos and 15 have knots. As in the original study, we classify a protein as exhibiting an age-associated structural change if at least one of its peptides exhibit a significant difference in abundance (defined as a |fold change| > 5 and adjusted p-value < 0.05) between aged and young cells^26^. 468 proteins exhibited a structural change, of which 443 have high-quality AlphaFold structures. We discard from our analyses those proteins containing knots and covalent lassos. After filtering, we have LiP-MS data for 2,256 proteins that have high-quality AlphaFold structures, including 1,686 that have one or more native NCLEs, and 438 proteins exhibited age-associated structural changes (Fig. 2a, see Dataset 1).

**Fig. 2:**
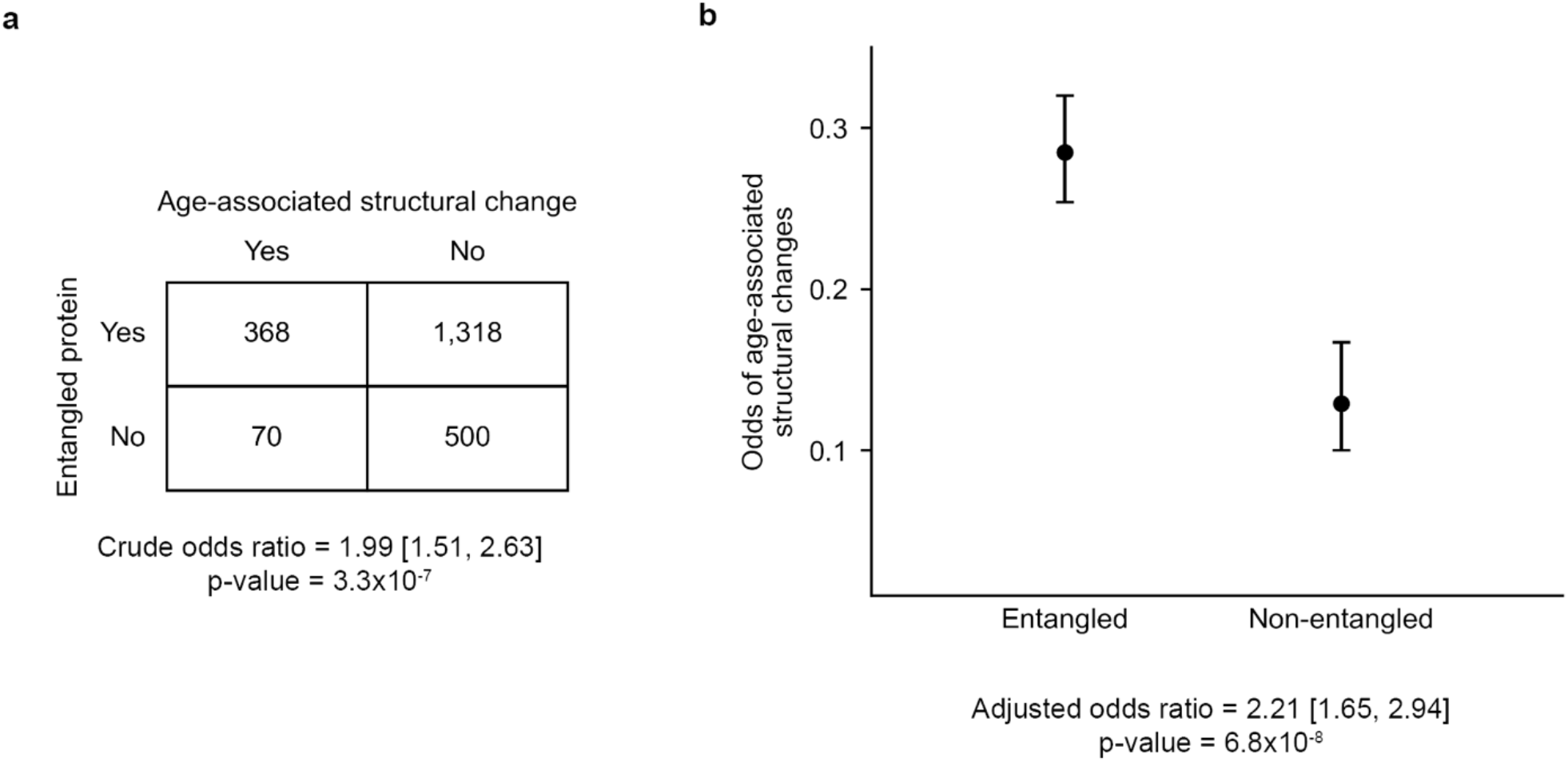
Association between the presence of native NCLEs and age-associated structural changes. (a) Contingency table and the crude odds ratio, summarizing the number of entangled and non-entangled proteins with or without age-associated structural changes. (b) Odds of age-associated structural changes between entangled and non-entangled proteins are estimated from the logistic regression model. Error bars represent 95% confidence intervals; adjusted odds ratio is estimated from logistic regression model.

### Natively entangled proteins are 121% more likely to exhibit an age-associated structural change

*E. coli* proteins with NCLEs are twice as likely to misfold upon chemically-induced unfolding and refolding as compared to those without these geometric motifs^22,28,29^. We hypothesized the same is true for *S. cerevisiae* proteins, and that this could cause the observed age-associated conformational changes. This hypothesis predicts that those proteins detected to exhibit such conformational changes by LiP-MS will be enriched in proteins containing native NCLEs. To test this prediction, we calculated both the crude odds ratio and the adjusted odds ratio which controls for confounding factors. (The odds ratio quantifies the odds of age-associated structural changes occurring in natively entangled proteins relative to natively non-entangled proteins.) For the former test we use a contingency table and the p-value is computed using Fisher’s exact test^30^, for the latter we used logistic regression^31^ to model the log odds of a protein exhibiting an age-associated structural change as a function of the presence of native NCLEs while accounting for protein length as a potential confounder since longer proteins are inherently more likely to misfold^32^ and contain native NCLEs^33^ (Methods).

Age-associated structural changes occur in 19.4% (=438 / 2,256 * 100%) of proteins across the set of mass-spectrometric-observed proteins. 21.8% (=368 / 1,686 * 100%) of proteins with one or more native NCLEs exhibit such a structural change, compared to 12.3% (=70 / 570 * 100%) among those without a native NCLE. The crude odds ratio of association between the presence of one or more native NCLEs and an age-associated structural changes is 1.99 (95% confidence interval (CI) = [1.51, 2.63], p-value = 3.3×10^−7^, Fisher’s exact test, Fig. 2a), indicating a positive association. The adjusted odds ratio (‘adjusted OR’) is 2.21 (95% CI = [1.65, 2.94]; p-value = 6.8×10^−8^, logistic regression controlling protein length as a confounder, Fig. 2b). We conclude that proteins with native entanglements are more likely to exhibit age-associated structural changes, increasing the odds by 121% (= (adjusted OR-1) *100%).

### Natively entangled regions are 59% more likely to exhibit age-associated structural changes

In *E. coli* proteins, entanglement misfolding is 40% more likely in those portions of the primary structure composing a NCLE compared to other portions of the protein^22^. Simulations indicated this was due to the greater likelihood of NCLEs segments being involved in misfolding events involving the failure-to-form the native NCLE^22^. We hypothesized the same is true for *S. cerevisiae* proteins, and that this could cause age-associated conformational changes to be localized in the natively entangled regions of a protein’s primary structure. Natively entangled regions comprise residues within ±3 positions along the primary sequence of the minimum loop-closing contact and crossing residue, as well as residues that are in contact with those residues composing the minimum loop-closing contact and crossing residues (see Fig. 3a, Methods). This hypothesis predicts that for the set of entangled proteins, the entangled regions will be more likely to exhibit age-related structural changes than non-entangled regions. To test this prediction, we again calculated the odds ratio using both contingency table and logistic regression to model the log odds of a protein residue exhibiting an age-associated structural change as a function of whether it composes a NCLE (Methods).

**Fig. 3:**
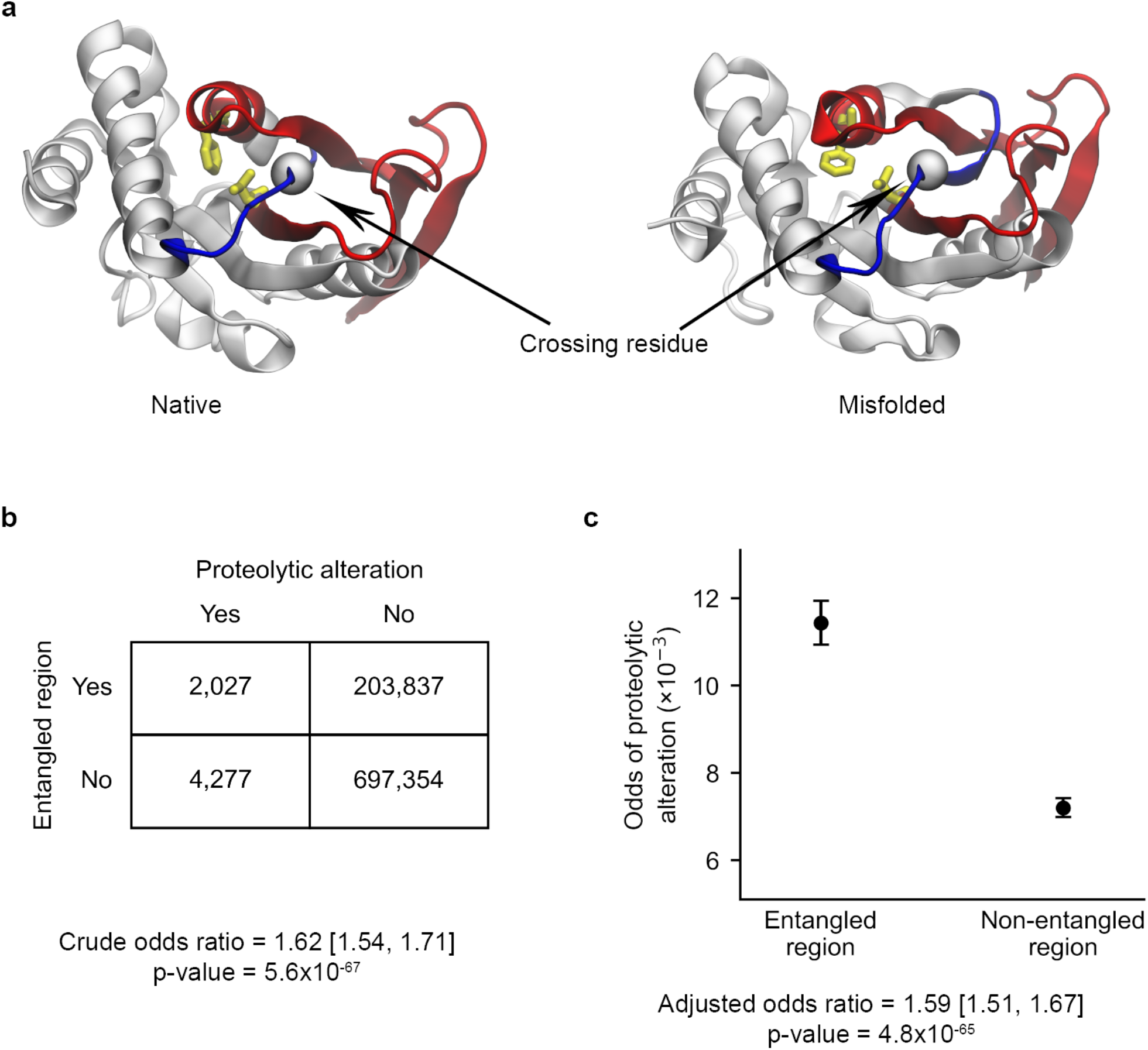
Enrichment of age-associated proteolytic alterations in entangled regions of natively entangled proteins. (a) Native and misfolded structures of adenine phosphoribosyltransferase 1 (Uniprot: P49435). The native structure contains a NCLE, whereas the misfolded structure – observed in simulations - exhibits loss of a native entanglement due to displacement of the threading segment outside the loop. NCLE components and color coding are consistent with Fig. 1a-b. (b) Contingency table and the crude odds ratio, summarizing the number of residues with or without age-associated proteolytic alterations in entangled and non-entangled regions. (c) Odds of proteolytic alteration between entangled and non-entangled regions are estimated from the logistic regression model. Error bars represent 95% confidence intervals; adjusted odds ratio is estimated from logistic regression model.

The prevalence of age-associated structural changes across the set of entangled protein residues is 0.70% (=6,304 / 907,495 * 100%). 0.99% (=2,027 / 205,864 * 100%) of residues composing entangled regions exhibit an age-associated structural change, compared to 0.61% (=4,277 / 701,631 * 100%) among the non-entangled residues (Fig. 3b). The crude odds ratio of association between being an entangled residue and exhibiting an age-associated structural change is 1.62 (95% CI = [1.54, 1.71], p-value = 5.6×10^−67^, Fisher’s exact test, Fig. 3b), indicating a positive association. The adjusted odds ratio is 1.59 (95% CI = [1.51, 1.67]; p-value = 4.8×10^-65^, logistic regression controlling protein length as a confounder, Fig. 3c). We conclude that not only are proteins with native entanglements more likely to exhibit age-related structural changes, but those changes are also more likely to occur in the regions of the protein involving a native entanglement, increasing the odds by 59% (=(adjusted OR-1) *100%).

### Natively entangled, age-associated proteins are seven times more likely to misfold

Why are age-related structural changes more frequent in entangled proteins and their entangled regions? Our hypothesis is that misfolding is more likely to occur in these proteins and occurs through a mechanism in which the native entanglement fails to form – that is, the loop closes but the threading segment never pierces it. This hypothesis predicts that between two similar groups of natively entangled and non-entangled proteins, there should be more misfolding in the former group. To test this prediction, we assembled two sets of proteins for simulation. The first set consists of 24 randomly selected proteins that contain native entanglements and exhibit age-associated structural changes in the LiP-MS data. The second set is pair-matched to have similar lengths to the first, but with no native NCLEs and exhibiting no age-associated structural changes (Methods, and Table S1). We then performed coarse-grained molecular dynamics refolding simulations by first thermally unfolding each protein and then temperature quenching them to 300 K to estimate their misfolding probability in the last 200 ns of 1.5 µs long trajectories (Eq. 8, Methods).

The proteins in the entangled and age-associated structural change group consistently showed greater populations of misfolded states (median = 27.9%, 95% CI = [12.0%, 45.2.%], bootstrap resampling of the median with 10^6^ iterations, *n*= 24 proteins) compared to the non-entangled and no age-associated structural change proteins (median = 4.1%, 95% CI = [0%, 20.1%], bootstrap resampling of the median with 10^6^ iterations, *n*= 24 proteins). This difference in medians is statistically significant (p-value = 2.5×10^-4^; one-sided Brunner-Munzel test^34^). Thus, the entangled and age-associated-structural-change proteins exhibit a sevenfold increase in misfolding probability compared to non-entangled and non-age-associated-structural-change proteins in this model (Fig. 4). We conclude that the age-associated structural changes in proteins could arise from the increased probability of natively entangled proteins to misfold.

**Fig. 4:**
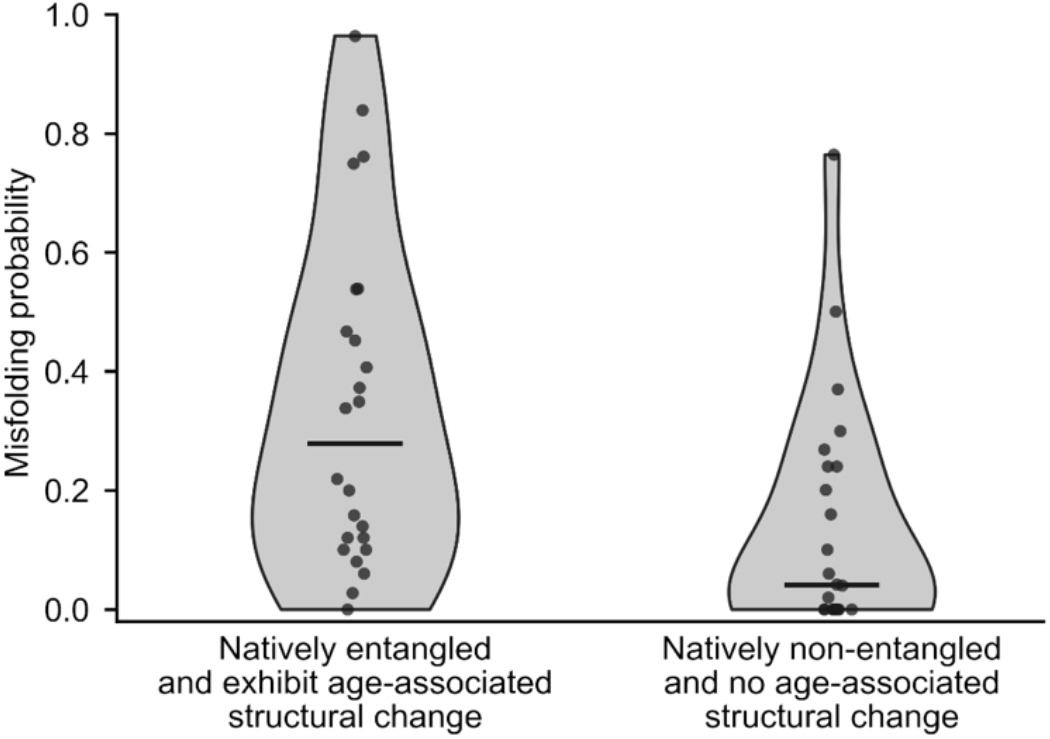
Misfolding probability from molecular simulations for entangled proteins with age-associated structural change and non-entangled proteins without age-associated structural change. Violin plots show the distribution of misfolding probability for proteins in each group (n = 24 proteins per group; proteins listed in Table S1). Points represent individual proteins, and horizontal lines indicate group medians.

### A structural basis for how misfolded proteins can accumulate with age

How entanglement-misfolded proteins can accumulate and persist during aging is a central mechanistic question. Previous studies indicate entanglement misfolded states can populate near native misfolded states that can evade the refolding action of chaperones^17^. We examine here if a similar mechanism might be possible for age-related structural change proteins. We focus on the GDP dissociation inhibitor GDI1 (YER136W; Uniprot: P39958), that contains 9 unique native entanglements and exhibits age-associated structural changes.

GDI1 is an essential cytosolic protein that regulates Rab GTPase recycling and vesicular trafficking in *S. cereviciea*^35^. We analysed the conformational ensemble of GDI1 obtained from our refolding simulations. Structural clustering reveals one state corresponding to the native state (state 6) and five distinct non-native states (Fig. 5a). In the last 200 ns of simulations, the native ensemble accounts for 62.8% of the population (Fig. 5b). Notably, states 4 and 5, which have an average fraction of native contacts *Q* value of around 0.9 (Eq. 6, Methods), are collectively populated 30% of the time. Thus, these two states are highly native like – 90% of their native contacts are formed – and are appreciably populated. This structural similarity is evident when a representative structure from state 4 is compared to the native structure (Fig. 5d). Thus, for this age-associated structural change protein, its misfolded state could plausibly be expected to bypass the proteostasis machinery to a similar extent as the native ensemble.

**Fig. 5:**
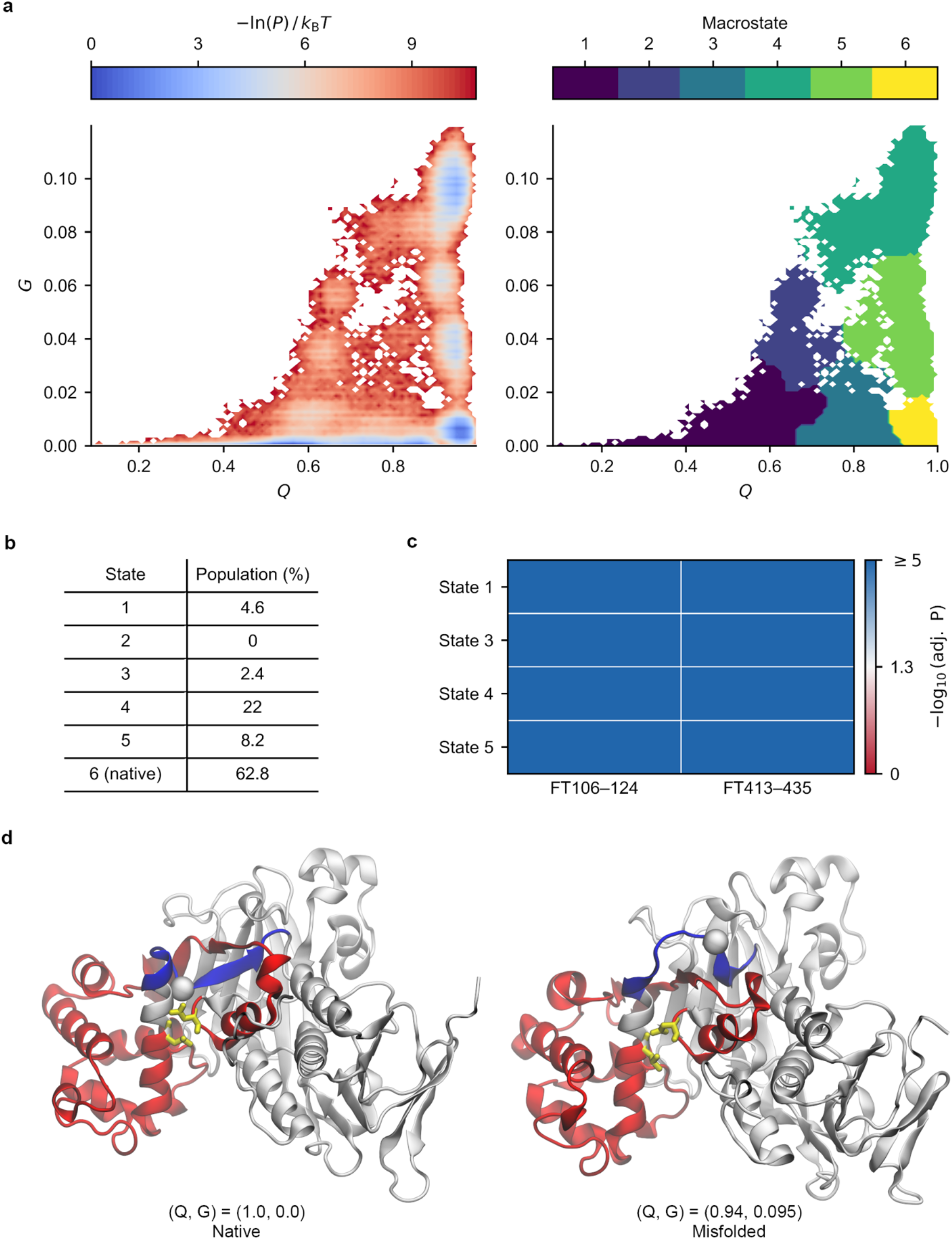
Entangled misfolded states are observed in simulations of GDP dissociation inhibitor GDI1. (a) −*ln*(*P*) surface and structural clusters identified from clustering analysis, in which *P* is the probability of a given state. (b) Population of the native and misfolded states evaluated over the final 200 ns of simulations. (c) -log_10_ (adjusted p-value) from permutation tests comparing the mean solvent accessible surface area of the proteolytically altered site (±5 residues) between each near-native misfolded state and the native state (state 6), ‘FT’ stands for full-tryptic peptide. (d) Representative structures of the native state (left, state 6) and a near-native misfolded state (right, state 4) illustrating loss of native entanglement. The entangled loop is shown in red and is closed by a native non-covalent contact between residues ASN92 and LEU219 (yellow licorice). The C-terminal threading segment is shown in blue. In the native state, the threading segment passes through the loop at the crossing residue PRO231 (white sphere), forming a non-covalent lasso entanglement. In the misfolded state, the threading segment is displaced outside the loop, resulting in loss of the native entanglement topology.

Next, we asked if the structures seen in our simulations are consistent with the alterations in proteolytic susceptibility detected by LiP-MS. To test this, we adopt the approach of a previous study^21^, in which proteolysis changes are assumed to arise from changes in solvent accessible surface area relative to the native state. In this approach, a metastable state is said to be consistent with the experimental data if the proteolytically altered region is also found to exhibit a significant deviation in surface area from the native state (Methods). We find that states 1, 3, 4, and 5 are all consistent with the age-related proteolytic changes (Fig. 5c, adjusted p − value < 0.05, or −log_10_ (adjusted p − value ) > 1.3 for these four states). This suggests they all have the potential to contribute to the observed digestion patterns, however, we conjecture that only the native-like states 4 and 5 will be relevant as the other states are more grossly misfolded and aggregate.

Taken together, these results suggest that a subpopulation of GDI1 occupies near-native misfolded states that structurally resemble the native ensemble but have a change in entanglement. Structures from these near-native misfolded states may persist for long-time scales since a substantial portion of the structure has properly folded into its stable native form. These misfolded states must unfold and backtrack to correct the misfolded topology, according to a previous study^21^.

### Proteins exhibiting age-associated structural changes are 52% more likely to increase in abundance with age

Misfolded proteins are typically less functional, and in aged cells with declining proteostasis capacity, may evade efficient clearance^12,16^ and accumulate^17^. We therefore hypothesized that proteins exhibiting age-associated structural change would be more likely to show increased abundance with age. To test this, we integrated LiP-MS–derived protein abundance measurements with transcriptomic (RNA-seq) profiling^26^. These protein abundances were normalized by their corresponding transcript levels to account for transcriptional differences between aged and young cells (Fig. 6a, Methods). Using logistic regression and controlling for protein length as a confounder, we found that proteins exhibiting age-associated structural changes are 52% more likely to increase in abundance with age compared to those not exhibiting age-associated structural changes (adjusted OR = 1.52, 95% CI = [1.22, 1.88], p-value = 1.5×10^-4^, logistic regression, Fig. 6b).

**Fig. 6:**
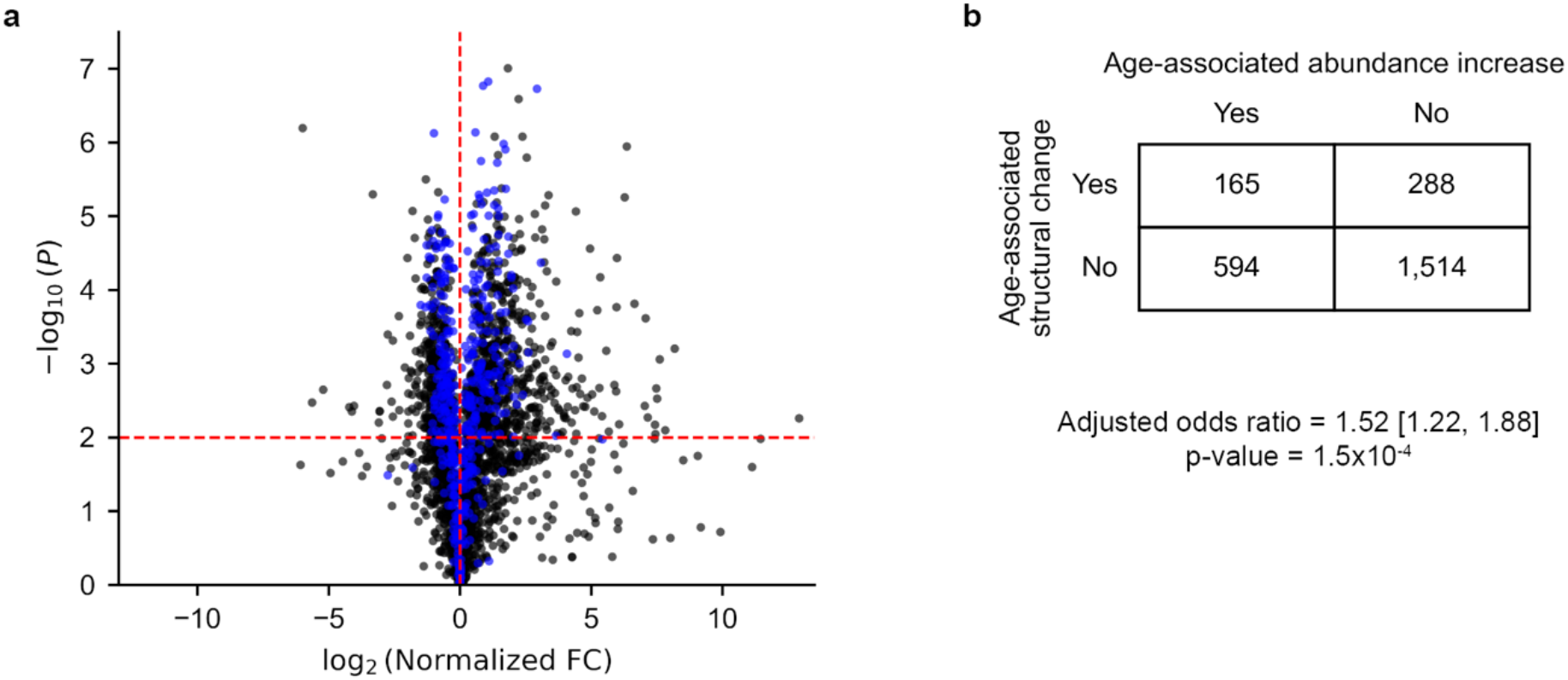
Age-associated structural change proteins are more likely to increase their abundance with age. (a) Volcano plot showing normalized log2 fold-changes in protein abundance between aged and young *S. cerevisiae* cells, after correcting for transcript-level changes. Each point represents a protein, with proteins exhibiting age-associated structural changes in blue and proteins not exhibiting age-associated structural changes in black. (b) Contingency table comparing the frequency of age-associated abundance increases in proteins exhibiting an age-associated structural changes versus not exhibiting age-associated structural changes.

Since these abundance changes were adjusted for differences in mRNA abundance with age, the observed increases in protein levels could result either from (1) increased translation initiation rates, where these proteins are translated more frequently in aged cells, or (2) reduced degradation efficiency of the misfolded states. The current data and analyses cannot differentiate between these two mechanisms. Further experiments, such as ribosome profiling would be required to do so. Regardless, we can conclude that these results are consistent with a model in which age-associated misfolding and loss of function are associated with increased protein abundance.

## Discussion

Our results indicate that native non-covalent lasso entanglements can give rise to a large proportion of the age-associated structural changes observed in the *S. cerevisiae* proteome. By integrating proteomics data from limited proteolysis mass spectrometry with entanglement annotations and molecular modelling, we found that proteins containing native NCLE motifs are twice as likely to exhibit age-associated structural changes than those without after four generations of yeast replication. These structural changes are not random across the protein’s structure but concentrated in the natively entangled segments of the protein that define the NCLE geometry – exhibiting a 59% higher likelihood of proteolytic alteration compared to non-entangled regions.

Our folding simulations provide insights into how native entanglements can contribute to these age-related structural changes. By simulating representative proteins that were both natively entangled and exhibited age-associated structural changes versus non-entangled proteins that did not exhibit age-associated structural changes proteins we observed a clear intrinsic effect of NCLEs on folding outcomes – the entangled proteins exhibited a seven-fold higher tendency to misfold compared to similarly sized, non-entangled proteins. These simulations were conducted under identical conditions, indicating that the difference in misfolding propensity is primarily due to the presence of NCLEs. Simulations indicate the NCLE creates a kinetic trap – the failure to form the native entanglement while the rest of the protein segments properly fold – that is an off-pathway intermediate in the folding landscape^19,21^. Proteins lacking native NCLEs are more likely to reach their native structure quickly because the failure-to-form mechanism is not possible for them. Together, these simulation results demonstrate that native NCLEs alone are sufficient to increase misfolding, providing a direct mechanistic link between protein geometry and age-associated structural changes. Indeed, these results are consistent with and reinforce a growing body of evidence that proteins with native NCLEs are at higher risk of misfolding and loss of function across a variety of organisms^18,22^.

In *S. cerevisiae* there are two established mechanisms by which a mother cell can accumulate misfolded protein with increasing replicative age. The first is asymmetric cell division^36–40^, in which an aging mother cell preferentially retains damaged or aggregated proteins, while the newborn daughter cell is relatively rejuvenated. As a result, misfolded proteins accumulate to ever higher concentrations over time. The second is proteostasis decline^9–12^ with age, which reduces the ability of the mother cell to remove misfolded proteins.

Our study suggests an additional mechanism may exist. In our detailed analysis of GDI1’s misfolded ensemble we found subpopulations with conformations that are highly native, with nearly all native contacts formed. The few native contacts that did not form exhibit a change in entanglement status, indicating the intertwining of backbone segments has localized, non-native arrangements. Recent studies have found that such near-native misfolded states, perhaps not surprisingly, can bypass the refolding actions of chaperones to a similar extent as natively folded proteins^17^. These results and observations suggest that these near-native misfolded states could accumulate by a failure of the proteostasis machinery to differentially act on them relative to the native state. Their continued presence in the cell can then further impair cellular functions.

In summary, our results suggest a new and novel mechanism of aging in cells. Near-native entanglement misfolding – primarily involving proteins with NCLEs in their native state – leads to soluble, non-functional conformations that cannot be efficiently cleared by the proteostasis network. As cells age, these misfolded states accumulate (Fig. 7) resulting in the structural changes detected by LiP-MS. This mechanism opens up new therapeutic targets and approaches to treat aging that focus on avoiding entanglement misfolding.

**Fig. 7:**
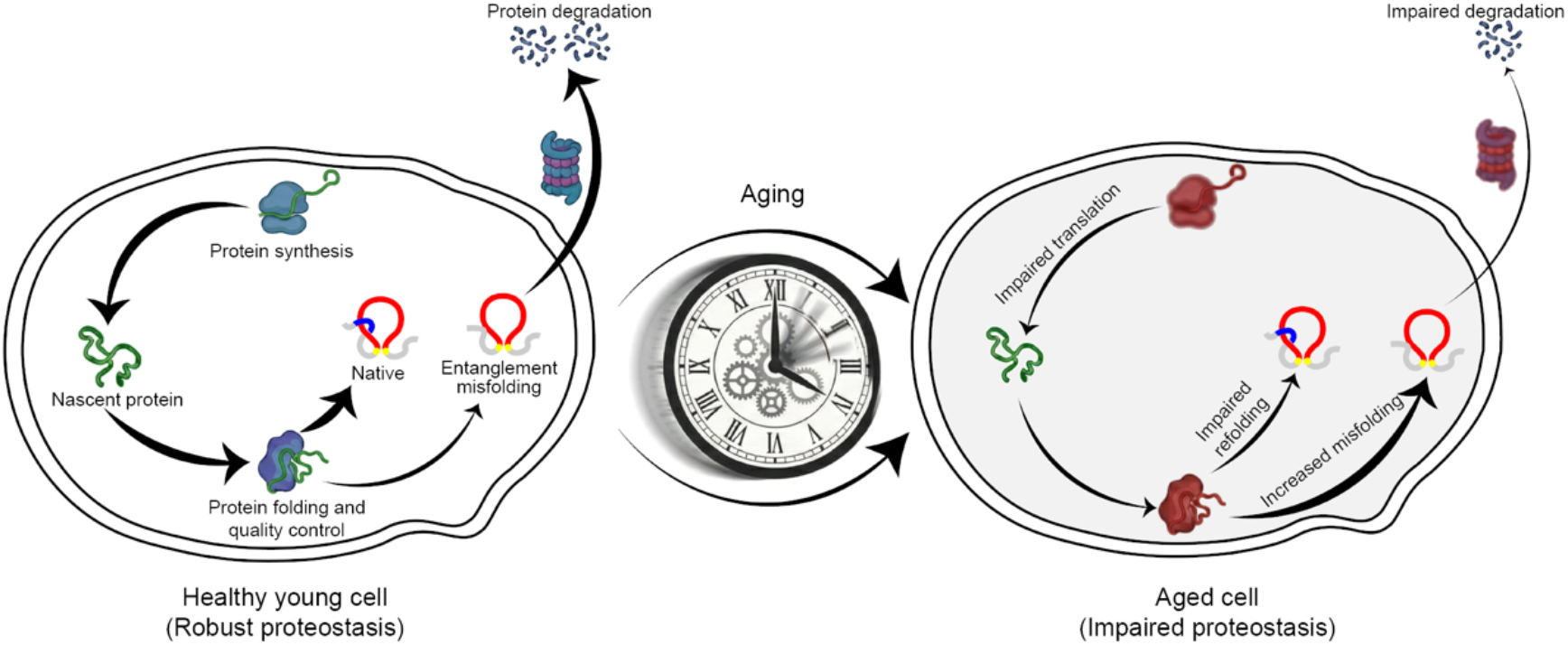
Entanglement misfolding contributes to age-associated structural changes. Proteins containing native NCLEs are prone to misfold into long-lived, near-native misfolded states. As cellular quality-control capacity declines with age, these misfolded states persist and accumulate, reducing protein function and contributing to widespread age-associated structural changes.

## Methods

### Construction of the dataset of proteins with native NCLEs and age-associated structural changes

To maximize structural coverage of the yeast proteome, we used AlphaFold2 predicted structures for *Saccharomyces cerevisiae* proteins^27^. We selected the structures with an average pLDDT score ≥ 70 to ensure that only proteins with reliable structural confidence were included in the analysis. Annotations of native NCLEs were obtained from a previously published dataset by our group^25^. In brief, the native NCLEs were identified by Gaussian Linking Integration method^18,41–43^. Two residues *i* and *j* are in contact that close the loop if they have any heavy atoms within 4.5 Å of each other. The linking numbers between N-terminus (*g*_N_) and the closed loop:

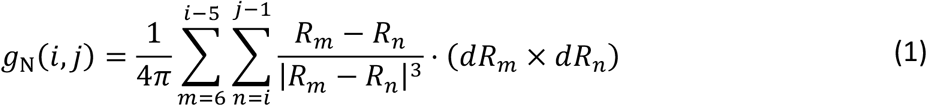

And the linking number between C-terminus (*g*_*c*_(*i,j*)) and the closed loop:

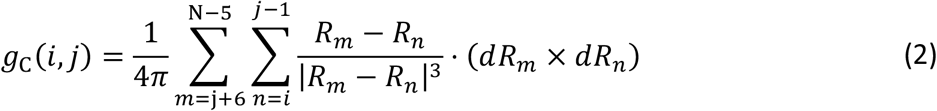

Where, ***R***_*l*_ and *d****R***_*l*_ are the average coordinates and the gradient of point *l* along the protein structure, were calculated as:

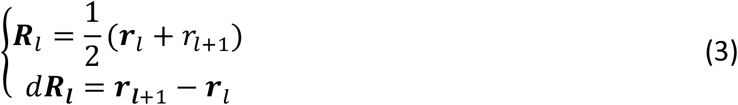

A given loop is determined to be entangled if |*g*_*N*_(*i,j*)| ≥ 0.6 or |*g*_*c*_(*i,j)*| ≥ 0.6. If an NCLE is detected for a given native contact, we use the Topoly^44^ Python package to identify the residues at which the termini cross the loop plane (crossing residues). Slipknots were excluded from our analyses. Because a single protein structure can contain multiple degenerate NCLEs (similar loop-closing contacts and crossing residues), we cluster them to obtain a set of unique representative NCLEs, as described in detail in our previous work^22,25^. Each protein was thus classified as natively entangled if it contains at least one NCLE in its predicted structure, or non-entangled if it does not have any NCLE.

The list of proteins exhibiting age-associated structural changes was taken from Paukštytė *et al*.^26^, which were identified from the yeast proteome-wide LiP-MS analysis on young (average age 0.6 ± 0.91 divisions) and aged (average age 4.2 ± 0.65 divisions) cells. A protein was classified as age-associated structural changes if it has at least one significant altered proteolytic cleavage mapped.

By intersecting the two datasets, we obtained a list of proteins that had both a high-confidence structure prediction and were observed in the LiP-MS aging study. Proteins containing knots or covalent lasso entanglement, as annotated by AlphaKnot 2.0^45^ and AlphaLasso^46^, were excluded from our analyses. This list formed the basis for our analyses linking native NCLE to age-associated structural changes.

### Modeling the association between native NCLE presence and age-associated structural changes

To evaluate the association between the presence of native NCLEs in proteins and age-associated structural changes, while control for covariate factors (e.g protein length), we employed a logistic regression model:

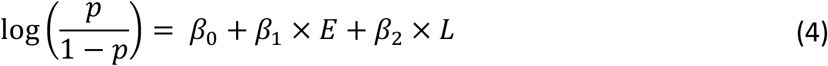

where *P* denotes the probability that a protein exhibits age-associated structural change (has at least one altered proteolytic peptide). *E* is a binary indicator of NCLE presence in protein native structure (1=entangled, 0=non-entangled), and *L* represents the standardized protein length, calculated by subtracting the mean and dividing by the standard deviation of protein lengths in the dataset. *β*_*i*_ (*i*=0,1,2) are the regression coefficients.

Protein length was included as a confounder to account for potential confounding, as longer proteins are intrinsically more likely to have altered peptides mapped to them^3^, and longer proteins are more likely to form NCLE conformation^33^. Logistic regression was performed using the statsmodels python package^47^.

### Modeling the enrichment of entangled regions in age-associated proteolytic alteration sites

To assess whether entangled regions are enriched for these age-associated proteolytic alterations sites, we employed an analogous logistic regression model to investigate, within entangled proteins, whether residues located in entangled regions are associated with altered proteolytic cleavage site between generation 4 and generation 0:

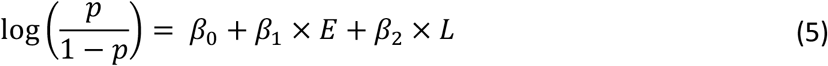

Here, *p*represents the probability that a residue exhibiting alteration of proteolytic cleavage. The altered proteolytic cleavage sites were derived from both half-tryptic and tryptic peptides that exhibit significant difference in abundance after normalizing for protein concentration between samples in generation 4 and generation 0. For half-tryptic peptides, we assigned the cleavage site to the terminus generated by proteinase K digestion. For tryptic peptides, the central residue was used to approximate the site of conformational perturbation, based on the assumption that structural changes within the peptide influence its overall accessibility. A buffer of ±3 residues along the primary sequence of altered proteolytic cleavage site was applied to account for potential positional uncertainty and structural fluctuations. *E* is the binary indicator specifying whether the residue is in the native entangled region. Native entangled region for each native representative NCLE were identified as follows: we first determined the loop-closing contact residues and the crossing residue(s) for each native representative NCLE. We then defined two residues sets. The first set included all residues within ±3 positions along the primary sequence of any loop-closing contact or crossing residue(s) of representative NCLE. The second set included any residue that formed at least one heavy-atom contact within 4.5 Å of loop-closing contact or the crossing residue(s). Finally, entangled region is the combination of these two sets. *L*represents the standardized protein length, calculated by subtracting the mean and dividing by the standard deviation of protein lengths in the dataset. *β*_*i*_ (*i*= 0,1,2) are the regression coefficients. Logistic regression was performed using the statsmodels python package^47^.

### Selection of protein candidates for coarse-grained simulations

To select protein candidates for coarse-grained simulations in the natively entangled and exhibit age-associated structural change group, we first identified all proteins containing native NCLEs that exhibited age-associated structural changes, had sequence lengths shorter than 800 residues, an asphericity parameter < 0.4, and a fraction of intrinsically disordered residues < 20% (estimated using MobiDB^48^). A total of 259 proteins satisfied these criteria. From this set, we randomly selected 20 proteins whose length distribution matched that of the original 259 entangled and age-associated structural change proteins (Fig. S1). In addition, we included four proteins: elongation factor 1-gamma 2, tRNA (guanine(26)-N(2))-dimethyltransferase mitochondrial, heat shock protein SSA1, and ATP-dependent molecular chaperone HSP82 — because of their known functional importance.

For each selected entangled age-associated structural change protein, we also identified a non-entangled and no age-associated structural change protein of comparable sequence length to serve as a control. The complete list of selected proteins is provided in Table S1.

### Coarse-grained modelling and temperature-quenching simulations

We use our previously published Cα coarse-grained representation with a Gō-based^49,50^ force field, in which each amino acid is represented by a single interaction site located at the Cα atom with a specific van der Waals radius.

The AlphaFold-predicted structure of each protein in our sets (natively entangled and age-associated structural change as well as natively non-entangled and no age-associated structural change) was first subjected to a high-temperature simulation at 600 K for 15 ns to generate unfolded conformations. The system was then cooled to 300 K to initialize refolding. All simulations were performed using a Langevin thermostat with a time step of 15 fs and a friction coefficient of 0.05 ps^-1^. For each protein, 50 independent trajectories were simulated, each lasting 1.5 µs. Mirror image trajectories were identified and removed using our previous protocol^19^, then replaced by new trajectory. Conformations were saved after every 5000 steps for analyses. Simulations were performed using OpenMM^51^ version 8.0.

### Calculation of fraction of native contacts *Q*and fraction of native contacts change in entanglement *G*

To characterize the nativeness of simulated conformations, we used the fraction of native contacts *Q* and the order parameter *G*, which measures the fraction of native contacts that exhibit a change in entanglement.

In the native structure of a protein, two residues are considered to form a native contact if they are separated by more than three residues along the primary sequence and have any pair of heavy atoms within 4.5 Å. For each frame of the simulation trajectory, the fraction of native contacts (*Q*) is computed as the ratio between the number of native contacts present in that frame and the total number of native contacts in the reference (native) structure. Given that we use a coarse-grained Cα model, a native contact is defined when the Cα–Cα distance between two residues in a frame is less than or equal to 1.2 times their corresponding Cα–Cα distance in the native structure:

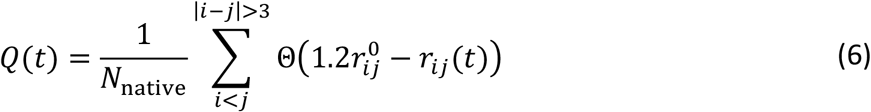

Where

*r*_*ij*_(*t*) is the Cα–Cα distance between residues *i*and *j*at time *t*,

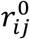is the corresponding native Cα–Cα distance,

*N*_native_ is the total number of native contacts in the reference structure, and

Θ(*x*) is the Heaviside step function (Θ(*x*) *=* 1if *x* ≥ 0, and 0 otherwise).

The fraction of native contacts with changes in entanglement (*G*) is also calculated per frame. A native contact is considered to exhibit a change in entanglement if its associated *g*_*N*_ or *g*_*c*_ value differs from that in the reference structure.

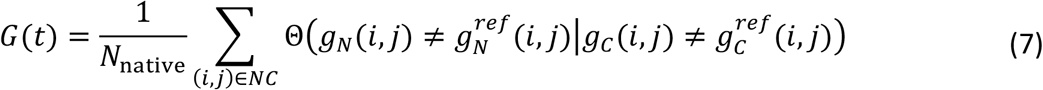

Where, NC is set of native contact in the reference structure.

### Structural clustering and misfolding probability estimation

The order parameters *Q* and *G* were used for structural clustering to categorize simulated conformations. The structural clustering was performed as follows: 1) The *Q* and *G* values were computed for each conformation in the simulation trajectory. 2) Each structure was assigned to a microstate using the Kmean++ clustering algorithm^52^, 100 microstates were used. 3) The resulting microstates were further grouped into macrostates using the PCCA+ algorithm^53^, number of macrostates was determined by visual inspection of the log-probability plot for each protein. This procedure yielded clusters of structures with similar degrees of nativeness (*Q*) and non-nativeness (*G*). All clustering analyses were carried out using the Deeptime package^54^.

To quantify misfolding, we defined misfolding probability for each protein as the fraction of near-native misfolded conformations sampled in the final 200 ns of the simulation trajectories. A structure is considered as near-native misfolded only if it belongs to a non-native cluster, according to clustering step. The misfolding probability *P*_mis_ for each protein is defined as:

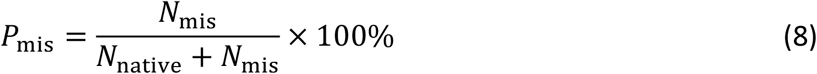

Where, *N*_mis_is the number of near-native misfolded conformations, and *N*_native_ is the number of native conformations in the last 200 ns of the trajectories.

### Normalization of protein abundance and identification of age-associated abundance increases

Protein abundance and transcript abundance measurements were obtained from the original experimental dataset referenced in this study^26^. For each protein, the mean abundance across three biological replicates was calculated for both young and aged samples. Protein abundance fold change was computed as the ratio of the mean abundance in aged cells relative to young cells.

To account for transcriptional changes during aging, protein fold changes were normalized by a transcript-derived normalization factor. Specifically, a normalization factor was calculated as the ratio of the mean transcript abundance across four biological replicates in aged versus young cells. Normalization was applied only to proteins whose transcript levels changed by at least two-fold in either direction (|log2 fold change| > 1) with p-value < 0.01, as determined by Welch’s t-test (two-tailed, unequal variance).

A protein was classified as exhibiting an age-associated abundance increase if its normalized fold change exceeded 1 (aged > young) and the corresponding Welch’s t-test yielded p-value < 0.01.

### Comparison of solvent accessibility of proteolytically altered regions between misfolded and native states

To evaluate the consistency between the misfolded ensembles from our simulations and experimental proteolysis data, we tested whether the solvent-accessible surface area (SASA) of regions near experimentally identified cleavage sites in the misfolded states are significantly different from those in the native ensemble. Specifically, for half-tryptic peptides, we calculated the SASA of residues surrounding the Proteinase K cleavage site, and for full-tryptic peptides, we used the central residue of each peptide as the reference point. In both cases, the SASA was computed for the region within 5 residues along the primary sequence around the selected site. The coarse-grained structures were first reconstructed using Pulchra^55^ to rebuild non-hydrogen atoms. Hydrogen atoms were then added in OpenMM using the Amber14 force field^56^. The resulting all-atom models were subjected to energy minimization for 100 iterations with harmonic restraints applied to the Cα atoms to preserve the global conformation of the coarse-grained models. Solvent accessibility for each proteolytically altered region was calculated using MDTraj^57^.

To determine whether the solvent accessibility of proteolytically altered regions in misfolded states are significantly different from that in the native state, we performed a numerical permutation test with 105 iterations. The test evaluated the null hypothesis that the ensemble means of SASA values for misfolded and native states are similar. The resulting p-values were adjusted using the Benjamini–Hochberg method^58^ to control the false discovery rate. If the adjusted p-value < 0.05, the null hypothesis was rejected, indicating a statistically significant difference in mean SASA between the misfolded and native ensembles.

## Data and code availability

All data supporting the findings of this study, along with the custom analysis code, are available on GitHub at (https://github.com/obrien-lab/entanglement_misfolding_and_aging)

## Acknowledgements

E.O. gratefully acknowledges support from the National Science Foundation National Synthesis Center for the Emergence of Molecular and Cellular Sciences NCEMS (DBI-2335029), as well as from the National Institutes of Health (R35-GM124818). Computational support was provided in part by Penn State Institute for Computational and Data Sciences (RRID:SCR_025154) and Pennsylvania State University RISE Core Facility (RRID:SCR_026426).

